# A simple DLP-bioprinting strategy produces cell-laden crypt-villous structures for an advanced 3D gut model

**DOI:** 10.1101/2022.02.09.479715

**Authors:** Núria Torras, Jon Zabalo, Eduardo Abril, Albane Carré, María García-Díaz, Elena Martínez

## Abstract

The intestine is a complex tissue with a characteristic three-dimensional (3D) crypt-villous architecture, which plays a key role in the intestinal function. This function is also regulated by the intestinal stroma that actively supports the intestinal epithelium, maintaining homeostasis. Efforts to account for the 3D complex structure of the intestinal tissue have been focused mainly in mimicking the epithelial barrier, while solutions to include the stromal compartment are scarce and unpractical to be used in routine experiments. Here we demonstrate that by employing an optimized bioink formulation and the suitable printing parameters it is possible to produce fibroblast-laden crypt-villous structures by means of digital light processing (DLP) stereolithography. This process provides excellent cell viability, accurate spatial resolution and high printing throughput, resulting in a robust biofabrication approach that yields functional gut mucosa tissues compatible with conventional testing techniques.

**Teaser:** 3D bioprinting approach for the direct fabrication of advanced cell-laden tissue constructs by means of visible-light photopolymerization.

## Introduction

*In vivo*, epithelial tissues are usually forming complex three-dimensional (3D) architectures that provide cells with specific microenvironments (1, 2). One of the most prominent examples is the small intestinal tissue, which folds forming finger-like protrusions called villi and invaginations called crypts. These microstructures are responsible for the exquisite cell type distribution along the crypt-villus axis of the tissue, the maintenance of cell renewal and homeostasis, the creation of oxygen gradients for the microbiome and maximizing absorption area and residence time for slow absorbing species (3–5). Indeed, it is now well recognized that *in vitro* models of intestinal tissue accounting for the 3D architecture provide barrier properties (permeability and transepithelial electrical resistance, TEER) that better resemble *in vivo* values, thus improving the predictability of *in vitro* assays (6, 7). However, it is also well known that the physiological function of the intestinal tissue does not only depend on a healthy epithelium, but also on the stromal tissue (named lamina propria) that lays below (2, 8). The entire structure, called intestinal mucosa, contains many cell types aside from the epithelial cells, including myofibroblasts, fibroblasts, endothelial cells and immune cells embedded in an extracellular matrix. It is therefore interesting to have access to intestinal tissue models that resemble not only the epithelium but the intestinal mucosa to properly model inflammatory bowel diseases, pathogen and microbiome interactions and even cancer (9, 10).

While in recent years there has been a lot of efforts to produce engineered tissues that recapitulate the architecture of the intestinal tissue, in most of the cases those have been limited to represent only the epithelium. Mostly, this is due to the use of complex fabrication methods based on replica moulding that are not friendly with embedding cells within the matrix (11, 12) or the use of relatively high dense matrices that do not allow cell survival (13, 14). Recently, some attempts have been made to include fibroblasts to mimic the stromal tissue in flat constructs (15), demonstrating the relevance of testing drug absorption in the presence of a full mucosa tissue. Also, there are some examples of including fibroblasts and other cell types in 3D structures (16, 17) employing replica moulding photolithography and laser ablation. These techniques, however, rely on the use of expensive equipment, and/or time-consuming procedures, which ends with a limited throughput. Therefore, the easy fabrication of cell friendly complex structures with soft materials (< 40 kPa) is still an open challenge.

Within this landscape, 3D bioprinting techniques offer unique features to tackle the problem. They offer a good resolution (compatible with most of the *in vivo* microstructures), good fabrication speed, they can be automatized and are cost-effective (18, 19). Among bioprinting, extrusion-based and light-based approaches have been employed to produce models of gut tissues (14, 20). Extrusion-based devices are robust systems compatible with standard culture plates. They use syringes, pistons, and screws to dispense drops or filaments of bioinks through microscale print heat nozzles (typically about 100 – 800 μm in diameter), which should be later stabilized in a postprocessing step using different triggers (e.g., temperature, light, or presence of ions) (21, 22). Combinations of alginate, gelatin, type-I collagen and decellularized extracellular matrix (dECM) powder have been proposed as bioinks for the fabrication of 3D printed matrices resembling the characteristic villous-like structures of the small intestine based on this approach (20, 23). Their main drawback of extrusion bioprinting is achieving continuous and free-form geometries with suitable resolution preserving shape fidelity, as well as achieving good survival rates, currently limited in the range of 40-86% due to the shear stresses inflicted on cells during the printing process (24, 25). Restrictions in bioinks’ viscosity and lower printing speeds are also major issues (26).

More recently, light-based bioprinting techniques have emerged as a strong alternative due to their low cost, simplicity in use and versatility. In particular, digital light processing (DLP) 3D bioprinting, based on stereolithographic (SLA) printing is gaining attention (27–29). DLP-SLA photopolymerizes layer-by layer bioinks, which can include a suspension of cells. These bioinks are located inside a cuvette (or vat) that has a transparent window at the bottom and allows for the projection of focused white and black patterns to create the 3D printed elements. 3D structures including protruded features and cavities can be replicated from CAD-based designs with high fidelity and a precise control on the layer thickness, achieving in-plane xy resolution up to 25 μm (30, 31). Due to the high versatility of the technique and the affordability of the system components, several customized DLP-SLA bioprinters have been reported so far, working with a variety of bioinks such as gelatin, hyaluronic acid (HA), poly (vinyl alcohol) (PVA) and polyethylene glycol (PEG)-based polymers combined with photosensitive elements (30–32), and light -sources (typically in UV and visible range) (33–35).

Most of the light-based bioprinting approaches rely on free-radical polymerization using acrylate-based polymers. Since this polymerization reaction is typically difficult to confine for low content and transparent polymer solutions, efforts have derived in strategies to gain control on pattern definition. For instance, PEG-based polymers have been successfully mixed with photoabsorbing species to improve the spatial control of the polymerization (30, 32). However, when targeting cell embedding with these materials, they lack cell-adhesion and degradation, impairing cell survival. An alternative is the use of natural derived hydrogels such as gelatin methacryloil (GelMA), which can also be crosslinked using light. Despite its good properties, when used at low concentrations, GelMA hydrogels are not mechanically stable, compromising the geometrical shape of the designs and lifespan of the printed constructs (36). Previous works have highlighted the benefits of combining PEG and GelMA polymers for the fabrication of cell-laden scaffolds for *in vitro* studies (13). Such combinations have also been successfully used as bioinks in 3D printing (37, 38).

Here we present a customized DLP-SLA 3D bioprinting system for the direct printing of tissue constructs using transparent, soft hydrogels, by means of visible-light photopolymerization. The system has been adapted to work with reduced bioink volumes and environmental control, allowing the fabrication of cell-laden structures resembling the intestinal mucosa in a single printing step. The intestinal tissues possess the 3D architecture of the small intestine, including villi and crypts and the epithelial and stromal compartments. Gelatin methacryloil (GelMA), poly(ethylene glycol) diacrylate (PEGDA) and visible-light lithium phenyl-2,4,6-trimethylbenzoylphosphinate (LAP) photoinitiator were used as main bioink components combined with photosensitive elements. With the proper tuning of (i) the bioink composition, (ii) the printing parameters such as layer thickness and exposure time, and (iii) the morphology of the CAD designs, we succeeded with the fabrication of 3D bioprinted intestinal tissues. Once printed, samples can be recovered with high fidelity and throughput, and allocated within standard Transwell^®^ inserts, where can sustain long-term culture with high cellular viability. Immunostaining analysis revealed cellular distribution and the expression of the main markers of both epithelial and stromal intestinal compartments, resembling the native intestinal tissue; proving the suitability of the technique for the fabrication of improved and more reliable engineered tissue models for *in vitro* studies.

## Results

### Bioprinting in a low-cost DLP-SLA apparatus

To successfully employ our low cost DLP-SLA 3D printing system as bioprinter, we customized the device including (i) an aluminium-based dedicated printing support and vat suitable for small solution volumes (< 2 mL), (ii) a flexible heater with a thermostat to keep the bioink solutions at 37°C allowing for the use of cell-laden polymers, and (iii) an infrared (IR) filter coupled to the output of the projector to minimize cell damage (Fig. 1A). The vat was equipped with an oxygen permeable transparent window made of fluorinated ethylene propylene (FEP) material. This allows the formation of a controlled liquid interface, known as “dead zone”, where molecular oxygen inhibits the crosslinking of the prepolymer solutions. This avoids the sticking of the newly formed hydrogel to the vat bottom surface and helps confining the photocrosslinked layer to the desired z plane (6, 28). For a given 3D design, the total printing time was adjusted by varying the total number of layers to be printed and their single exposure times. It should be noticed that reduced exposure times may cause weak crosslinking, whereas increased layer thicknesses will result detrimental for *z* resolution.

**Fig. 1:**
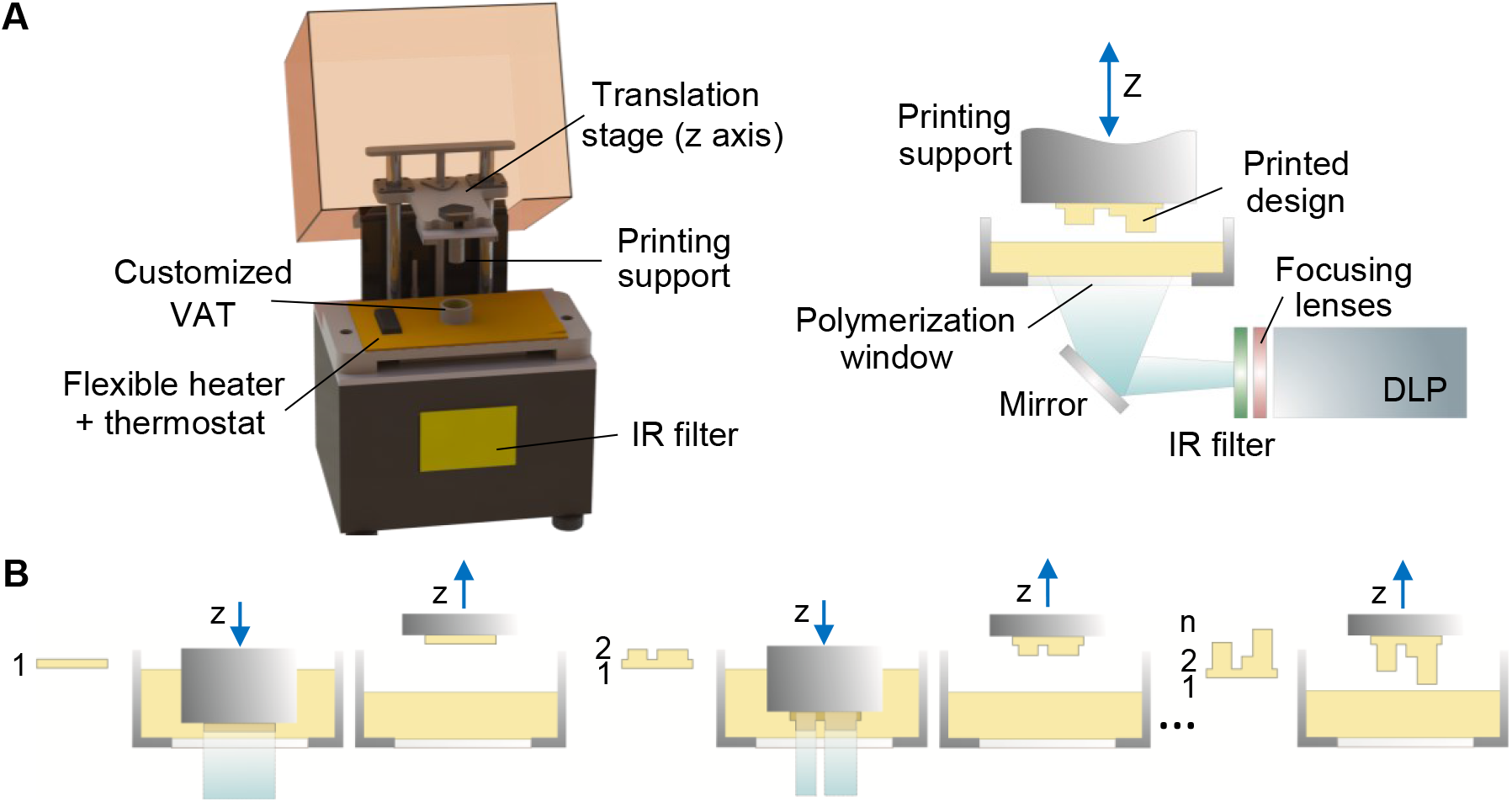
Custom 3D bioprinting system: (A) 3D CAD model of our 3D bioprinter system (left) with detailed view of the main components (right). (B) Scheme of the operation principle, which is based on layer-by-layer photopolymerization. Series of consecutive prints with different patterns result in final printed design.

The photocrosslinked layer thickness, also called cured depth (*C_d_*), depends on the effective energy dose to which the bioink is exposed. On the irradiated surface, the energy dose (E) depends on the power of the light source and the exposure time. However, light is absorbed as it passes through the prepolymer solution. Jacob’s equation, a semi-empirical equation derived from the Beer-Lambert law, is used to describe this exponential decay in light intensity, providing a relationship between the cured depth, the surface energy dose, the optical penetration depth (*D_p_*), and the critical energy to induce polymerization (Ec) (see eq.1).

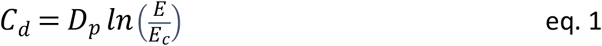

From this expression it can be inferred that a minimum energy dose is required to reach polymerization (Fig. 2A). On the other hand, *D_p_* is related with the composition of the bioink and strongly depends on the photoinitiator used, being both its molar extinction coefficient (ε) and its concentration, (χ) the most relevant parameters involved (see eq.2) (29).

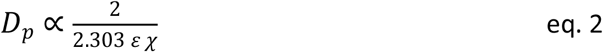

**Fig. 2:**
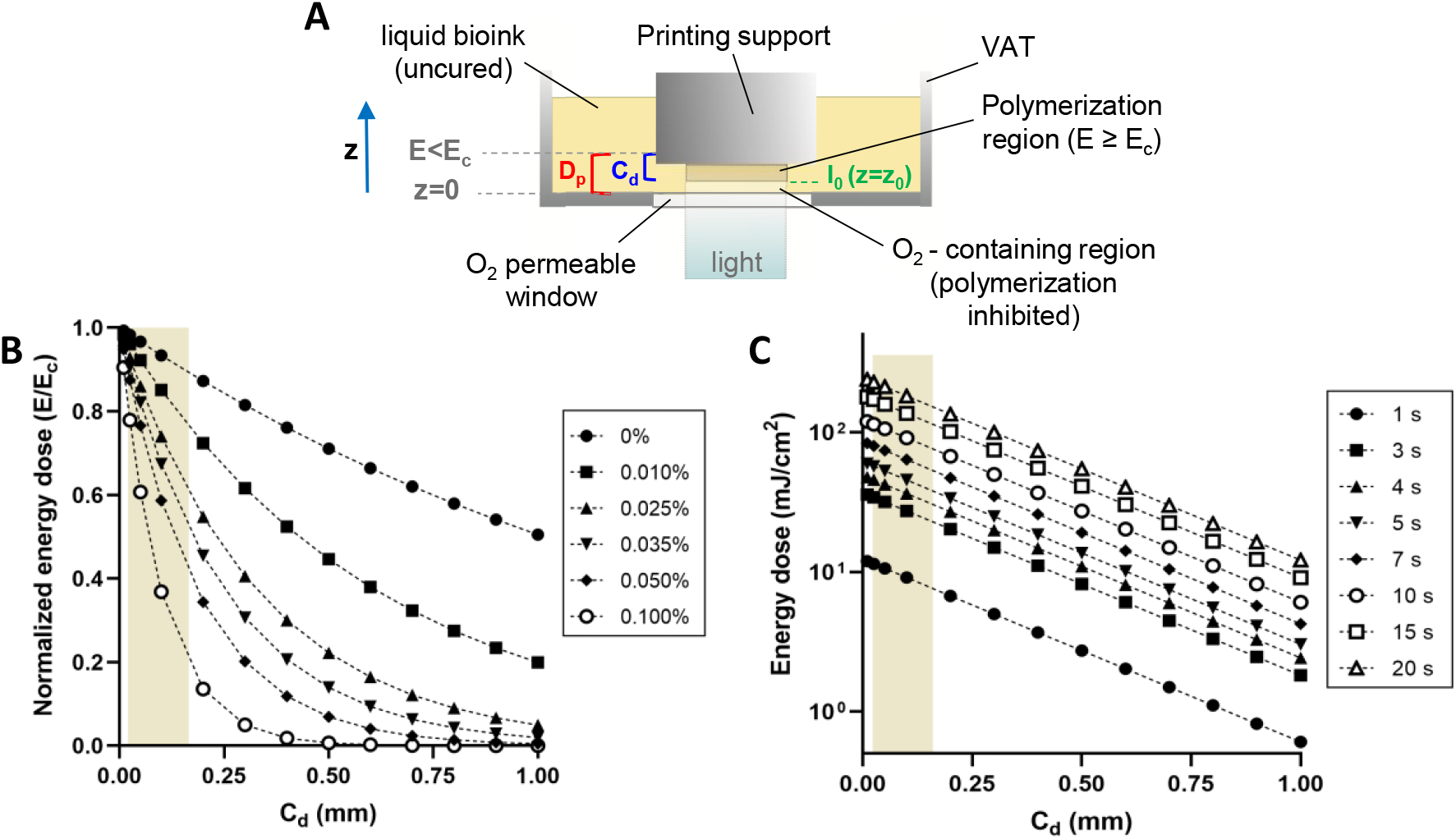
Theoretical analysis of the influence of the main printing parameters on the curing depth. (A) Sketch of the printing process showing the main elements involved in the photocrosslinking reaction. Effects of (B) the photoabsorbing components in the bioink composition and (C) the single layer exposure time on the resulting curing depth, *C_d_*. Colored region denotes the experimental working range.

In addition, the presence of molecular oxygen and other chemical species, such as photoabsorbers, photosensitive dyes or light-attenuating additives, has an impact on the photopolymerization reaction (29, 39). These species will inhibit the photopolymerization reaction or will absorb light, helping to confine the polymerization to the desired layer thickness and boosting the printing resolution (32). Therefore, Dp can be tuned varying the concentration of the photoabsorbing species in the bioink to predict the suitability of a photocrosslinkable solution to be printed. In our case, we used a bioink formulation based on GelMA-PEGA hydrogel co-networks, which has been proved to be printable (32, 40) and a suitable candidate for tissue engineering applications (13, 41). While PEGDA provides mechanical stability to the printed structures, GelMA offers good cellular response favouring their growth and attachment. In particular, our bioink of interest contained 5% w/v of GelMA and 3% w/v of PEGDA, together with 0.4 % w/v of visible-light LAP photoinitiator. To improve the resolution of our printing results, an additional photoabsorbing agent, tartrazine, was added to the bioink mixture. Tartrazine is a synthetic azo dye highly soluble in water at low concentration, with an absorption peak centred around 436 nm (42). Its absorbance overlaps the emission spectrum of our light source, and at the same time, it is close to LAP absorption peak (~375 nm), resulting in a beneficial mixture for printing well defined 3D structures (see Fig. S1). By varying the concentration of tartrazine in the bioink, the effects on the crosslinking and the shape and size of the resulting prints (*i.e*. the effects on the curing depth Cd for our bioink) could be estimated (Fig. 2B). Thus, for our particular bioink formulation, Cd expression can be rewritten as

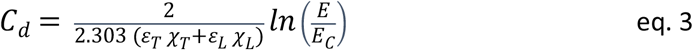

were *ε_T_* and *ε_L_* are the molar extinction coefficients of both tartrazine and LAP photoinitiatior, and *χ_T_* and *χ_L_*, their concentrations, respectively.

Considering the previous expression and the optical power density values at the crosslinking starting point, *z_0_*, (see Fig. S2), it is possible to predict the effect of the exposure time on the resulting Cd, for a given bioink formulation (Fig. 2C) and thus, anticipate the range of energy dose required for reaching a proper crosslinking and the maximum gel thickness reached (43).

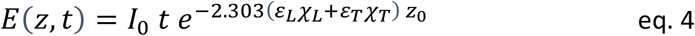

### Effects of the main printing parameters on the morphology of villous-like scaffolds

To check the influence of the main printing parameters (concentration of photoabsorbing agent, single layer exposure time and layer thickness) (Table 1), on the morphology of the printed constructs, 3D scaffolds mimicking the villous morphology of the intestinal tissue were produced. For this purpose, CAD designs featuring pillar structures of 700 μm in heigh and 300 μm in diameter spaced 750 μm on a 6.0 mm diameter and 150 μm thick disc-like base (see scheme in Fig. 3A) were printed and inspected following the procedure described in materials and methods section.

**Fig. 3:**
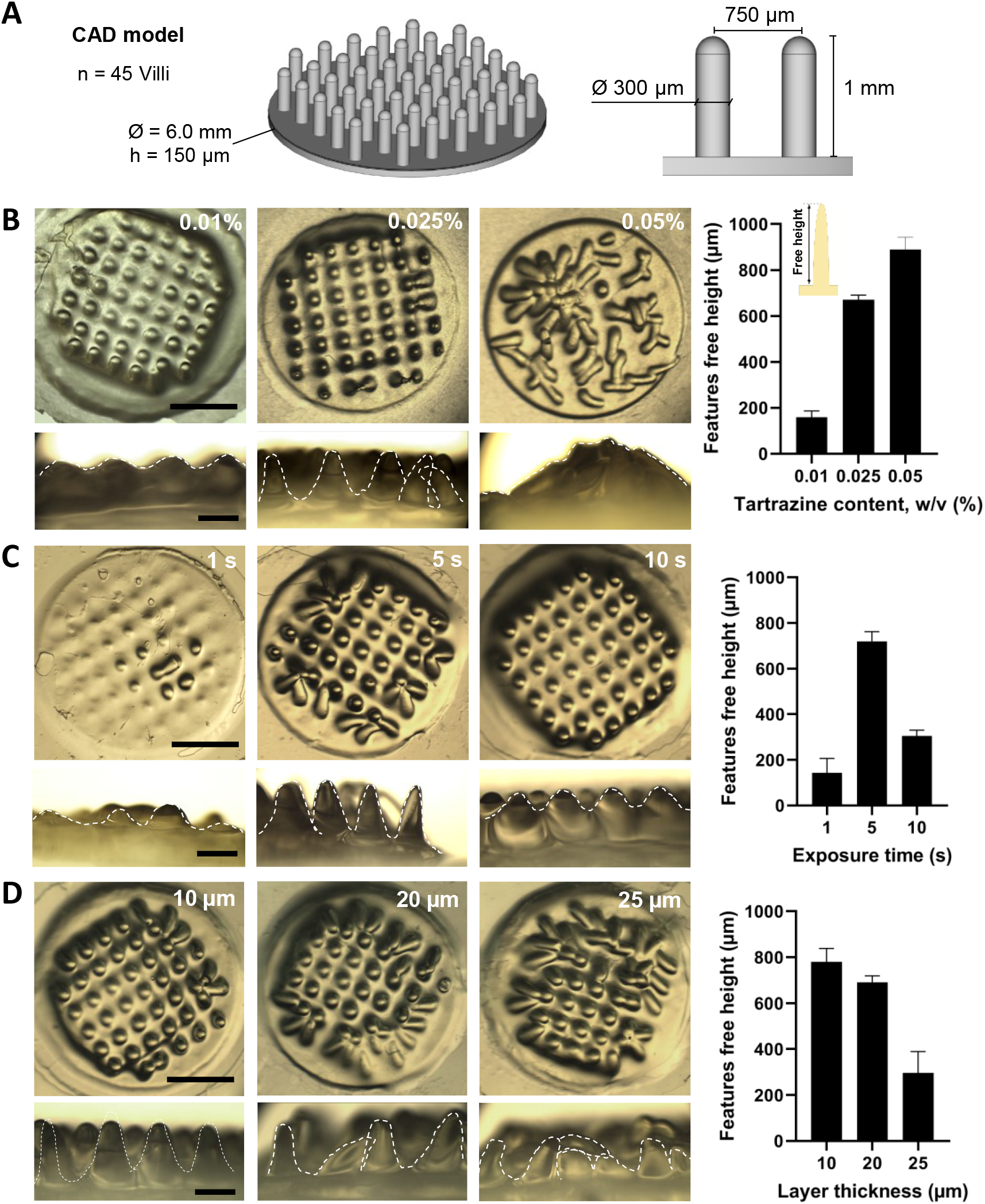
Effect of the main printing parameters on the features definition and free height. (A) CAD design with the main pillar dimensions used to study the effect of the printing parameters on the final prints. (B-D) Top and lateral views of 3D printed villous-like scaffolds fabricated with 5% w/v GeLMA and 3% w/v PEGDA copolymer with 0.4% w/v of LAP (left), and pillars free height quantification (right), as function of the tartrazine concentration in the bioink (B), the exposure time used per printed layer (C), and the layer thickness defined (D). Scale bars = 2 mm (top views) and 500 μm (lateral views).

**Table 1:** Values for the different printing parameters tested on the final prints

| Printing parameters | Values tested |
| --- | --- |
| Photoabsorber content (% w/v) | [0 - 0.075] |
| Exposure time per layer (s) | [1 - 15] |
| Layer thickness ($\mu\text{m}$) | [10 - 25] |

#### • Effects of the concentration of the photoabsorbing agent on the morphology of villous-like scaffolds

By taking into account the theoretical relationship between the different parameters shown in the previous section and thus the existing correlation between the bioink composition, the light propagation and the crosslinking kinetics, the effects of the photoabsorbent concentration on the morphology of the villi and crypts printed features were evaluated, fixing the printing parameters to 61.5 mJ/cm^2^ as exposure energy dose per layer, with a predefined thickness of 13 μm (net z motor displacement). Then, we added tartrazine at three different concentrations, 0.01, 0.025 and 0.05 % w/v (0.19, 0.47 and 0.94 mM, respectively), to the prepolymer GelMA-PEGDA-LAP mixture. According to Fig. 2B, all these concentrations, which are below the cytotoxic range of this dye (2.5 - 4 mM) (44, 45), should result in different values of optical penetration depth, *D_p_*. As observed in Fig. 3B, for the lowest tartrazine concentration (left panels), the crosslinking confinement was not efficient and it was extended to the regions with no direct light exposure, so the resulting structures appeared overexposed. As a consequence, villous-like protrusions looked as partially embedded within a thick base, resulting in a significant decrease on their free height (tip to base distance) of about 80%, compared to the intended dimensions (Fig. 3B). When increasing the tartrazine concentration to 0.025% w/v the crosslinking reaction was better confined, and significantly thinner and larger features were obtained for the same printing parameters (Fig. 3B, middle panels). For the highest tartrazine concentration tested (0.05% w/v), the crosslinking was confined very efficiently, and the features were well defined but, for the CAD design employed, the finger-like structures had such a high aspect ratio that did not self-stand (Fig. 3B, right panels). By the addition of tartrazine, the global molar extinction coefficient of the bioink increases up to 7.8-fold for the concentration of 0.05% w/v (ε_LAP_ = 218 M^-1^cm^-1^, at 365 nm; ε_Tartrazine_ = 21600 M^-1^cm^-1^, at 430 nm), resulting in dramatic decrease of the light penetration depth, *D_p_*, from 3.37 mm when there is no tartrazine to 0.43 mm for the highest tartrazine concentration tested (see eq. 2). In addition, as the effective energy diminishes, there is also a dose-dependent delay in the induction of the photocrosslinking, which affects the gelation kinetics of the hydrogel and might also help to confine the crosslinking reaction (32). In here we took advantage of this phenomenon to minimize undesired overexposure effects and fine-tune the morphology of the printing features.

#### • Effects of the single layer exposure time on the morphology of the villous-like scaffolds

To check the effects of the exposure time per layer on the morphology of the villous-like scaffolds, a bioink containing of the previous GelMA-PEGDA-LAP prepolymer mixture together with 0.025% w/v of tartrazine was used, applying an incident optical power of 12.3 mW/cm^2^ (Fig. S2) and a thickness per printed layer of 18 μm. The total exposure energy dose (defined as the incident optical power applied per exposure time) per layer was controlled by varying the single layer exposure time between 1 and 10 seconds. This provided a theoretical surface energy dose that, according to eq. 1, ranged between 6.67 and 86.21 mJ/cm^2^ for our experimental window, which defines a curing depth between 0.025 and 0.150 mm (Fig. 2C). A poorly crosslinked hydrogel was obtained for the lowest exposure time tested (Fig. 3C, left panels), probably because the energy dose applied per printed layer E (12.3 mJ/cm^2^) was lower to the estimated critical energy needed for the polymer chains to achieve enough crosslinking degree and form a gel, Ec (~35 mJ/cm^2^) for our particular approach. Increasing the exposure time up to 5 seconds, the total exposure energy dose per single layer significantly increased (see Fig. 2C), producing nice finger-like structures of a free height matching the one defined by the CAD design and with the proper aspect ratio (Fig. 3C, middle panels). On the contrary, longer exposure times (10 seconds), with surface energy dose E ≈ 104.24 mJ/cm^2^ increase the curing depth and allow the propagation of the crosslinking reaction to regions not illuminated by the pattern, again triggering an overexposure reaction (Fig. 3C, right panel). Through these experiments one can infer that, for a given pre-polymer solution, the crosslinking degree and confinement can be tuned by the selecting the proper energy dose of individual layers in a narrow range of time exposures to fine shape the morphology of the printed structures.

#### • Effects of the layer thickness on the morphology of villous-like scaffolds

In DLP-SLA systems, the total printing time (also known as building time) is dependent on the predefined layer thickness (*i.e., Cd*) and the single layer exposure time (29). So, in terms of throughput, large layer thicknesses and short exposure times should be beneficial. However, in the previous section it has been demonstrated that exposure times that are too short turned into poorly crosslinked hydrogels. Here, to check the impact of the layer thickness on the morphology of the villous-like scaffolds, we fixed the GelMA-PEGDA-LAP-tartrazine bioink mixture (5:3:0.4:0.025, all in % w/v), 61.5 mJ/cm^2^ as exposure energy dose per layer (12.3 mW/cm^2^ of incident optical power and 5 s of single exposure time), varying the layer thickness between 10 and 25 μm. As shown in Fig. 3D (left panel), the smallest thickness values per printed layer tested resulted in self-standing villous-like features, with some overexposure localized at the surrounding regions, but nice free feature heights close to those imposed by the CAD design. Thicker layer thicknesses, however, resulted in less defined features that did not self-stand (Fig. 3D, middle and right panels). These results are in agreement with previous findings above and proved that, for a given bioink composition and energy dosage, lower voxel units (*i.e*. shorter layer thicknesses) can be achieved, enhancing the printing resolution (30).

#### • Effects of the CAD design on the morphology of crypt-villous scaffolds

As our goal was to mimic the small intestine architecture for cell culture experiments, different CAD models including both extruded features and holes resembling the villi and the crypts of the tissue, respectively, were produced and printed. Crypts and villi heights, diameters, and distances among them (*i.e*., pitch) were varied according to reported values for this tissue in humans (Fig. 4A) (46). A hydrogel base in the shape of a thin disc was added as handling support for the whole structure as well as for the cells to feel the mechanical properties of the soft material. Fig. 4B shows three examples of different CAD models (left) with their corresponding printing results (right), using the GelMA-PEGDA-LAP-tartrazine bioink mixture previously optimized, 61.5 mJ/cm^2^ as exposure energy dose and 13 μm of thickness per printed layer.

**Fig. 4:**
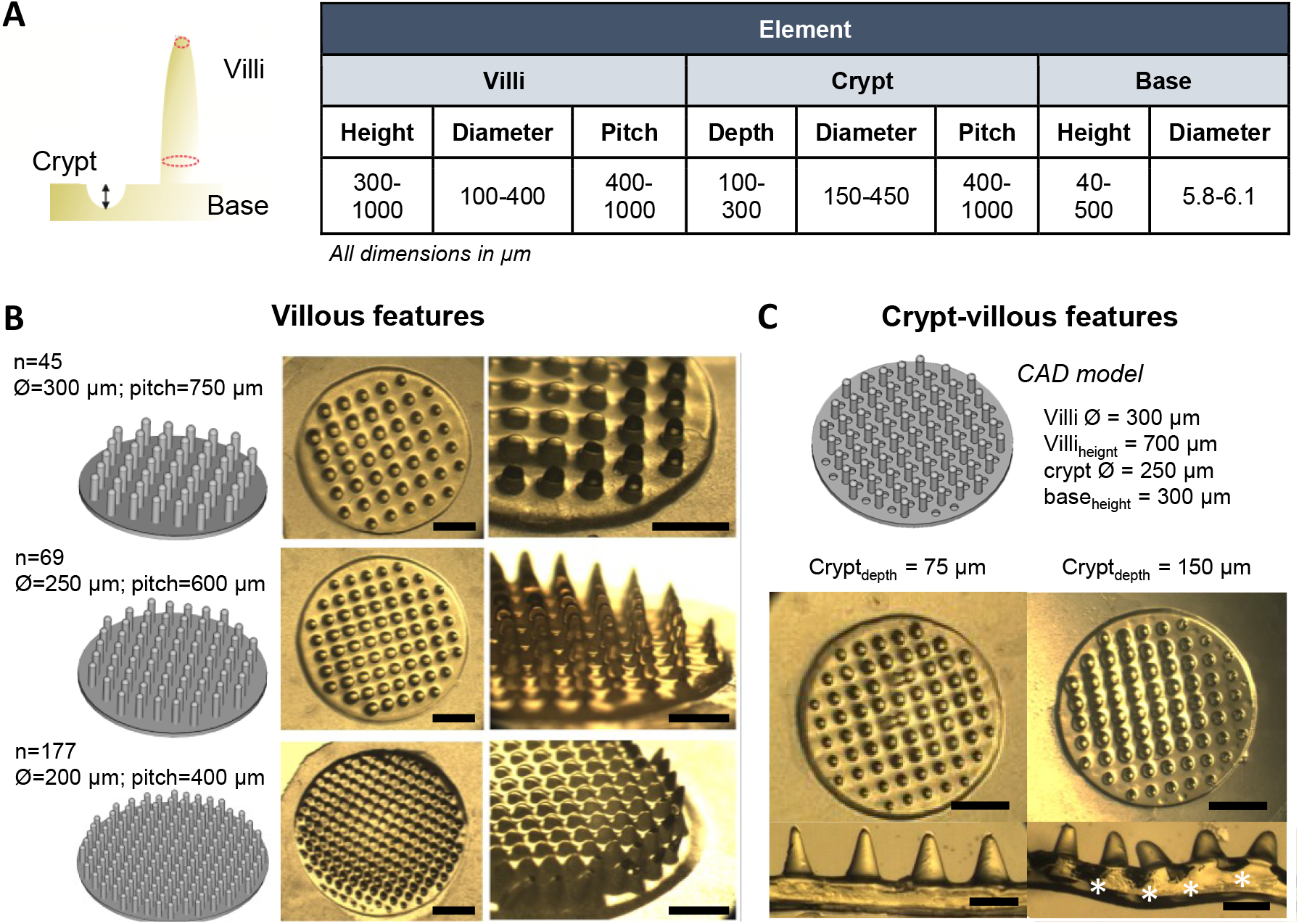
Effects of the CAD model dimensions on the morphology of crypt-villous scaffolds. (A) Range of the values tested for the different parameters in the CAD models mimicking the main intestinal features. (B-C) Examples of different CAD models and the corresponding printing results obtained using 5% w/v GeLMA, 3% w/v PEGDA, 0.4% w/v LAP and 0.025% v/v tartrazine as bioink. (B) Effect of the variations in distance between consecutive pillars: CAD models (left) and top (middle) and lateral (right) views of the printed results. Scale bars = 1.5 mm (top views), 1.0 mm (side views). (C) Effect of the crypt depth: CAD model (top) and top view (middle) and cross-sections (bottom) of the printed scaffolds, showing absence (left) or presence (right) of crypts. Scale bars = 1.5 mm (top views), 400 μm (cross-sections).

As it can be observed, there is a close relationship between the diameter of the villous-like features and the pitch between them (distance between centres). When this relationship is 2:5 or higher, prints with homogeneous, well-defined self-standing features can be obtained (Fig. 4B, middle panel). However, when this ratio is reduced to 1:2 or lower, polymer crosslinking takes place in between pillars (Fig. 4B, bottom). A similar effect is produced when crypt features are included in the CAD design (Fig. 4C). Experimentally, we observed that, within the range of dimensions tested (Fig. 4A, table), if the ratio between the crypts’ depth and their diameter is 1:3 or lower, crypts cannot not be distinguished, neither from the top nor within the cross-sections of the printed structure (Fig. 4C, bottom left). However, when this relationship is closer or larger than 3:5, crypts homogeneous in depth are fully visible both on the surface of the prints and in the cross-sections (Fig. 4C, bottom right).

By integrating the knowledge generated with the printing experiments performed, an optimal concentration of tartrazine of 0.025% w/v was selected to add to the bioink composed of 5% w/v GelMA, 3% w/v PEGDA and 0.4% w/v LAP. For this bioink composition, we selected as main printing parameters 13 μm of layer thickness with 5 s of exposure time per layer (61.5 mJ/cm^2^ of exposure energy dose) and we printed 3D structures with a base thickness of 250 μm, invaginations of 150 μm, and protrusions of 700 μm in a single bioprinting process, leading to crypt-villous scaffolds with physiologically relevant dimensions.

### Swelling, mechanical and diffusion properties of bioprinted GelMA-PEGDA hydrogels of optimized composition

The network architecture of hydrogels is directly correlated with their water uptake capacity, which affects the penetration and transport of essential nutrients and oxygen (47). Hence, it is relevant to characterize the swelling properties of the hydrogels produced using the bioink formulation and printing parameters leading to the optimized crypt-villous scaffolds. Swelling properties were quantitatively assessed by measuring the volumetric swelling ratio, SV of the hydrogels. GelMA-PEGDA-LAP-tartrazine disc-like hydrogels were printed using the previously optimized parameters and allowed to reach equilibrium by incubation in HBSS buffer at 37°C for 5 days. At predefined timepoints, the discs were imaged, and their diameters and heights were measured to obtain S_V_ and the variation along the radial, S_R_ and transversal, S_T_, directions (Fig. 5A). Initially, samples experienced an abrupt increase of their volume within the first 30 minutes describing a peak of 15.2% +/- 0.8% (S_Vmax_), that rapidly decreased and stabilized forming a plateau, resulting in S_V_ = 11.6% +/- 2.0% at equilibrium after 24 h. Sample dimensions along both radial and transversal directions (Fig. 5B and 5C) followed a similar trend, with a more pronounced variation in the sample height.

**Fig. 5:**
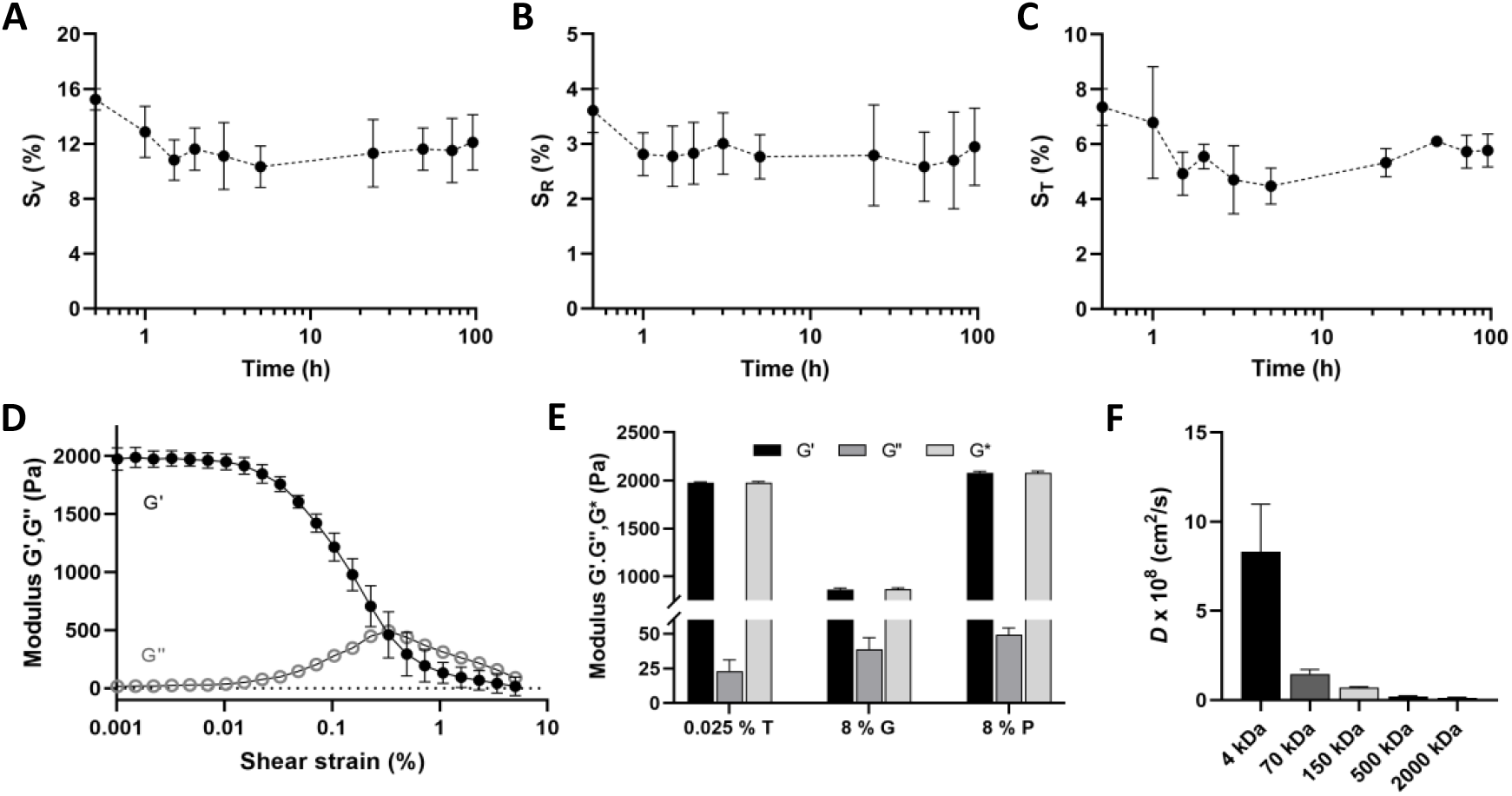
Swelling, mechanical and diffusive properties of the bioprinted hydrogels: (A) Volumetric swelling measurements of disc-like samples during 96 h. (B-C) Swelling ratios on radial, SR, and transversal, ST, sample directions. (D-E) Mechanical performance of the printed gels for samples containing 0.025% w/v of photoabsorber. Rheological curves showing G’ and G’’ moduli (D) and comparison of complex, storage and loss moduli (E). (F) Diffusion coefficients for dextran molecules of different sizes diffusing through the bioprinted hydrogels (FD2000 = 200 kDa, with a hydrodynamic diameter of 41.6 nm; FD500 = 500 kDa, with a hydrodynamic diameter of 32 nm; FD150 = 150 kDa, with a hydrodynamic diameter of 17 nm; FD70 = 70 kDa, hydrodynamic diameter of 11.6 nm; FD4 = 4 kDa, hydrodynamic diameter of 2.8 nm). Results are shown as mean ± SD (min n=3).

The swelling ratio measured for our 3D bioprinted hydrogels was significantly lower than previous values reported for GelMA-PEGDA co-networks of similar macromer content (3.75% w/v GelMA - 3.75% w/v PEGDA) but different polymer rate (13). It is recognized that GelMA-PEGDA hydrogels exhibit tailorable swelling properties, which primary depend on the GelMA’s methacrylation degree and the total macromer concentration. PEGDA is a hydrophilic polymer whose molecules have two binding sides at the end of the chain, whereas GelMA ones have multiple crosslinking points all over the chain, in agreement with its methacrylation degree. Thus, increasing the GelMA content of our bioink, denser networks can be achieved, leading to lower swelling rates due to its decreased interaction with water molecules (47). Furthermore, the printing parameters chosen, such as the energy dose applied, and the nature and concentration of the photoinitiator used, have also been proved to have a direct impact on the swelling behaviour of the bioink (48). Additionally, in this case, the presence of a photoabsorbing agent in the bioink composition affects the photopolymerization process (Fig. 3B), also altering the swelling properties.

In addition to swelling, another parameter that is of paramount importance when embedding cells in a 3D matrix is the stiffness of such matrix, which ideally should be close to the *in vivo* tissue. For the intestinal tissue, elastic moduli values ranging from 1 to 100 kPa are reported (49, 50). Fig. 5D shows the components of the complex shear modulus measured as a function of the shear strain. The plots revealed a typical behaviour characteristic of viscoelastic solids (G’ > G’’) with a certain degree of crosslinking in its network structure (51). We also determined that, as previously observed in the printing results and the swelling behaviour, the crosslinking degree and thus, the mechanical properties, can be tuned by just changing the tartrazine concentration (Fig. S6). For the bioink solution employed, which includes tartrazine in a concentration of 0.025 % w/v, the storage modulus G’, measured from the linear region of the curve (up to 0.01% of the shear stress) was 2.07 ± 0.41 kPa. This value lays between those obtained for 8% w/v GelMA and 8% w/v PEGDA, being closer to the latest (Fig. 5E). Considering our hydrogels as quasi-isotropic materials (Poisson’s coefficient of ~0.5), the apparent elastic moduli when adding 0.025 % w/v of tartrazine to the bioink composition resulted in 5.94 ± 0.19 kPa, evidencing the softness of our gels. Moreover, these values obtained in agreement with others reported in literature for PEGDA- and GelMA-based hydrogels (6, 52, 53), as well as with the ones from other materials typically employed for *in vitro* 3D tissue cultures, such as Matrigel^®^ and Collagen I, measured using similar standard assays (54). In addition, the mechanical properties measured for our bioprinted hydrogels are within the range of the ones described for *in vivo* soft tissues (49).

3D matrices for cell cultures should also guarantee the transfer of nutrients and oxygen and the excretion of waste metabolites. These mass transfer properties are related with the porous structure of the bioprinted GelMA-PEGDA hydrogels. To gain insight on that, the diffusion profiles of model dextran compounds of different molecular sizes were measured, and their diffusion coefficients through the hydrogel co-network were estimated (Fig. 5F). While the hydrogel acted as a diffusion barrier for large molecules (500 and 2000 kDa, see Fig. S4), it allowed the diffusion of small (4 kDa) and medium size molecules (70 and 150 kDa). The size exclusion limit, defined as the smallest dextran diameter excluded from the pores, was determined from the plot of the diffusion coefficients versus the *In* (MW) (see Fig. S4) (55). The molecular weight exclusion limit was found to be 360 kDa (215 to 750 kDa, 95% CI). This pore size ensures good mass transfer properties for the cell survival within the hydrogels since it allows for diffusion of molecules such as albumin (58 kDa), one of the most abundant proteins present in cell culture media.

### GelMA-PEGDA hydrogels bioprinted with the low cost DLP-SLP instrument sustain cell culture effectively

NIH-3T3 mouse embryonic fibroblasts were chosen to validate the suitability of our setup for bioprinting, since they are commonly used for cell encapsulation experiments (13, 56). In addition, these cells represent the fibroblast cell population within the stromal compartment of our intestinal model. For this purpose, 7.5·10^6^ cells/mL were mixed with the optimized prepolymer solution containing 5% w/v GelMA, 3% w/v PEGDA, 0.4% w/v LAP photoinitiator and 0.025% w/v tartrazine dissolved in HBSS supplemented with 1% v/v Penicillin/Streptomycin to form the bioink. The mixture was placed in the vat, a simple squared-based CAD design containing a 3×3 grid was loaded, and the printing process was carried out using the above optimized printing parameters (see Fig. 6A). Cell-laden samples were printed on top of treated porous PET membranes to be later mounted on Transwell^®^ inserts for cell culture. Fig. 6B shows bright field images of a bioprinted grid-like structure containing the embedded NIH-3T3 cells.

**Fig. 6:**
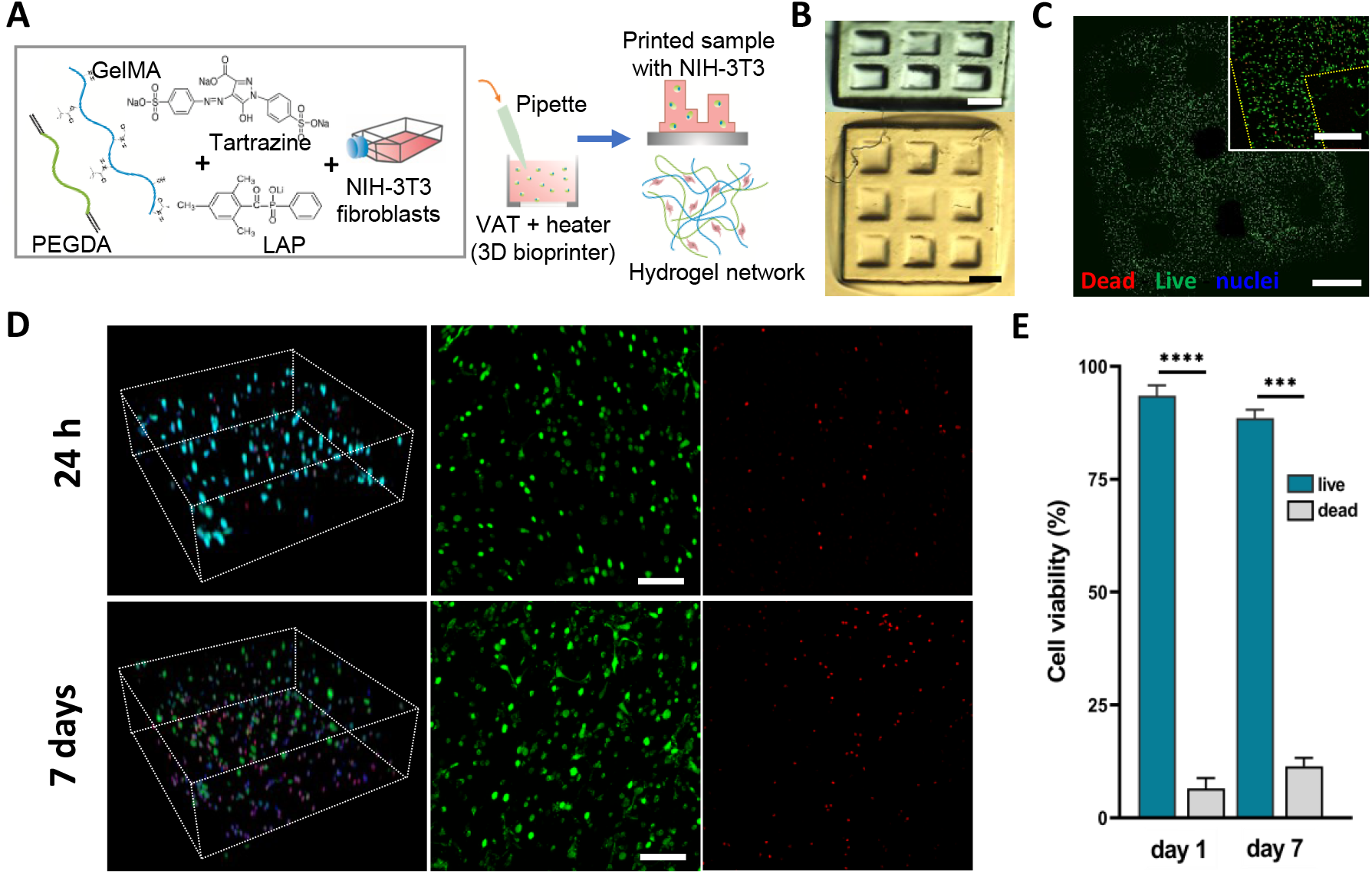
Validation as bioprinting system: (A) Schematic to illustrate the procedure followed for NIH-3T3 fibroblast encapsulation in hydrogel co-networks. (B) Bright field images of a 3D printed grid-like structure. Scale bars = 1 mm. (C) NIH-3T3 cell-laden 3D bioprinted grid at 24 h after live/dead assay (live cells stain in green, dead cells in red). Hoechst was used to stain the nuclei. The image was constructed by stitching tiled fluorescent images. The inset shows a detailed view of one corner, where cells were starting to spread. Scale bars = 1 mm and 400 μm, respectively. (D) Confocal 3D reconstruction (left) and maximum intensity projections (right) of samples at 24 h and 7 days after encapsulation. Scale bars = 100 μm. (E) Cell viability quantification based on live/dead staining. Values are the mean percentage of cell viability +/- SD (n=4). **** p<0.0001; *** p=0.008.

Cell viability within the bioprinted hydrogels was investigated using a Calcein-AM/ethidium homodimer live/dead assay and confocal microscopy 24 hours and 7 days after cell encapsulation. After 24 hours, most of cells were alive and homogeneously distributed within the whole volume of the hydrogel; being the ones closer to the surface and corners starting to spread (Fig. 6C). Overall, the cell viability after the bioprinting process was excellent with this method (above 93%, Figs. 6D, 6E). This might be related to a friendly bioprinting process and to the relatively small swelling of the bioinks we used, which minimizes cell stress after printing (13). In addition, after 7 days in culture, cells appeared evenly distributed within the hydrogels, being most of them spread. Cell viability slightly decreased with respect to the values obtained 24 hours after encapsulation, but still remained high, above 86% (Figs. 6D, 6E). These results proved that both the bioink composition selected and the printing parameters, result in hydrogels able to sustain cell culture in an effective manner.

### Cell-laden GelMA-PEGDA bioprinted hydrogels sustain a functional 3D *in vitro* model of the small intestinal mucosa

Once proved the capability of our system for creating suitable 3D microstructured environments for cell culture, we moved a step forward combining cell-laden GelMA-PEGDA bioprints with the co-culture of epithelial cells on top, thus creating a 3D *in vitro* model of the small intestinal mucosa.

Fibroblast-laden hydrogels including villous and crypt features were bioprinted mixing GelMA-PEGDA bioink with NIH-3T3 cells at a density of 7.5·10^6^ cells/mL (Fig. 7A). After the bioprinting process, samples were mounted in Transwell^®^ inserts and 3.5·10^5^ Caco-2 cells were seeded on top (Fig. 7Ba). Samples were kept in culture for 21 days, monitoring the cell growth and monolayer formation. Fig. 7Bb shows top view brightfield pictures of the Caco-2 cells grown on top of fibroblast-laden hydrogels and cell-free hydrogels, used as controls, at different culture time points. 1 day after seeding, most of the Caco-2 cells both in the co-cultured and control samples remained clustered on the base of the 3D structures, covering the crypts. After 11 days, samples in co-culture were fully covered by an epithelial monolayer, whereas in controls some villous tips and base regions remained uncovered (Fig. 7Bb middle). After 21 days of culture and contrary to the controls, the co-cultured samples maintained intact epithelial monolayers. Along the culture time, the effective barrier function of the epithelial monolayers was assessed by measuring the transepithelial electrical resistance (TEER) (Fig. 7Bc). Consistent with the microscopy observations, control samples showed very low TEER values for all the time points tested, corresponding to uncomplete epithelial monolayers. In contrast, TEER values of fibroblast-laden hydrogels with Caco-2 cells on top experienced a boost after 9-11 days in culture and increase until forming a plateau from day 15 on, reaching values of ~ 200 Ω cm^2^. These values are well in agreement with those reported for Caco-2 epithelial monolayers grown on 3D villous structures (6). Therefore, from these experiments we can conclude that the presence of the fibroblasts favours the growth of the epithelial monolayers on top of GelMA-PEGDA bioprinted hydrogels. These findings are in agreement with previous works that reported increased epithelial proliferation and differentiation in co-cultures of fibroblasts and epithelial cells (13, 56, 57). Moreover, we hypothesize that the boost in TEER values measured from days 9-11 of culture may likely match the time that the fibroblasts need to spread, migrate, and secret extracellular matrix proteins that contribute to the epithelial basement membrane. To explore this hypothesis, we then proceeded to perform immunostainings on histological cuts.

**Fig. 7:**
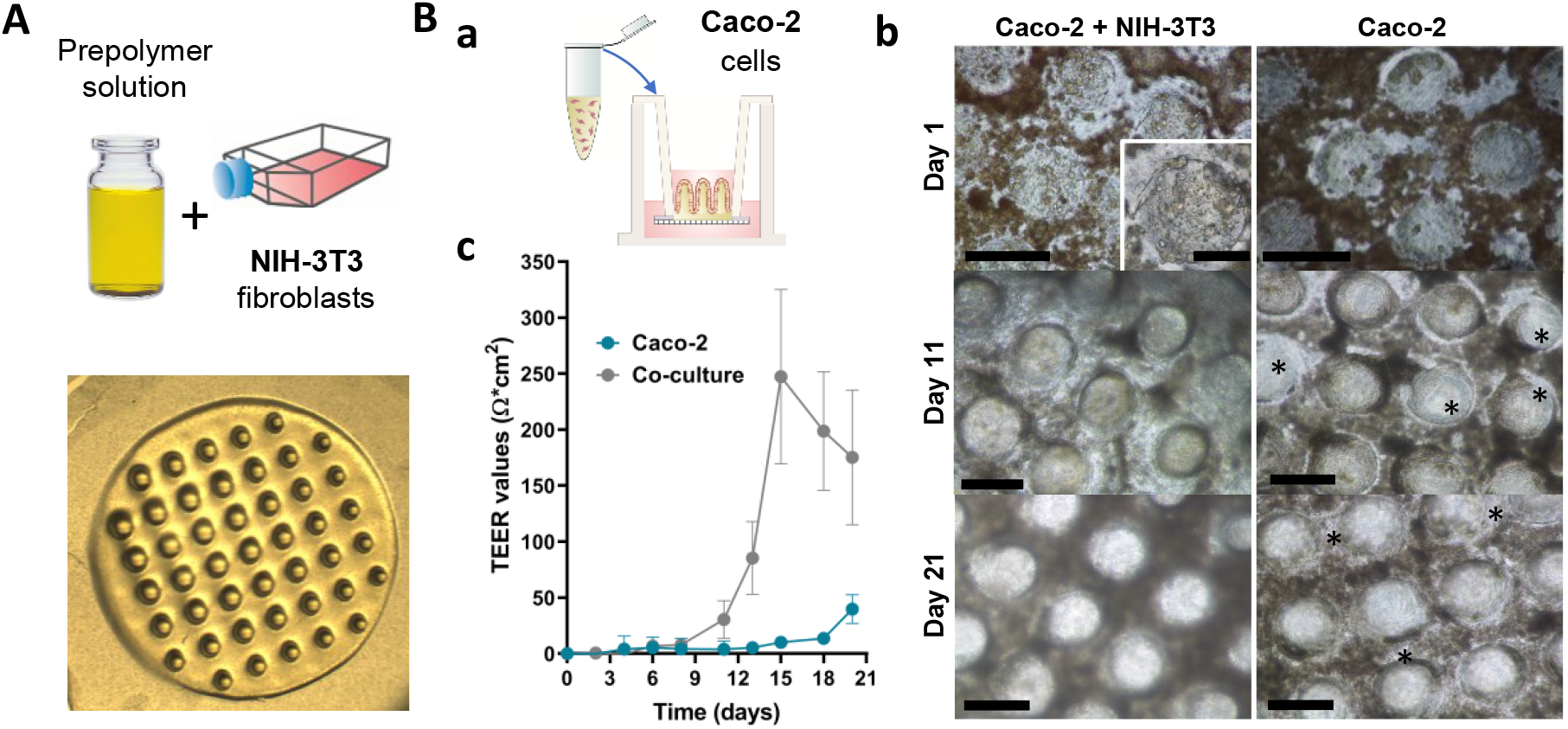
Monitoring of the 3D printed intestinal mucosa model: (A) Schematic illustration of the bioink composition and brightfield picture showing the printed model. Scale bar = 1.5 mm. (B) Schematic illustration of the Caco-2 seeding process on printed structures previously assembled on modified Transwell^®^ insert (a). Brightfield images (b) and TEER values (c) showing the Caco-2 monolayer progression on top of the printed hydrogels with (right-left) and without (right-right) containing embedded NIH-3T3. * denote regions non-covered by the epithelial monolayer. Scale bars = 100 μm, 50 μm (inset).

After 21 days in co-culture, samples were removed from the Transwell^®^ inserts and fixed to proceed with histological cuts. To protect the integrity of the soft 3D bioprinted structures, samples were first embedded in low molecular weight PEGDA (575 kDa), following the dedicated protocol developed by our group (58). Thereafter, samples were embedded in OCT^®^ medium and cross-sectioned before immunostaining and imaging.

Confocal images of sample histological sections revealed the formation of Caco-2 cell monolayers that, in agreement with the brightfield pictures and TEER measurements, fully covered the 3D surfaces in the co-culture samples. Whereas Caco-2 cell monolayers grown on the control samples showed discontinuities, mainly at the crypt regions (Fig. 8A). Epithelial cells grown on top of both fibroblast-laden and control hydrogels showed accumulation of F-actin at their apical region, which is compatible with the presence of brush borders in polarized epithelia. Supporting proper cell polarization, immunostainings also revealed the expression of β-catenin and ZO-1 markers at their proper locations for polarized epithelium (Fig. 8B). In addition, epithelial morphology appeared as evolving from columnar at the crypts and the tips of the villi, to cuboidal along their walls for both samples (Fig. 8B inserts). Concomitantly, epithelial cell density also varied along the 3D structures. These findings agree with the results found in similar 3D *in vitro* models (58) and, more importantly, with results reported from *in vivo* studies the intestinal epithelium (59, 60).

**Fig. 8:**
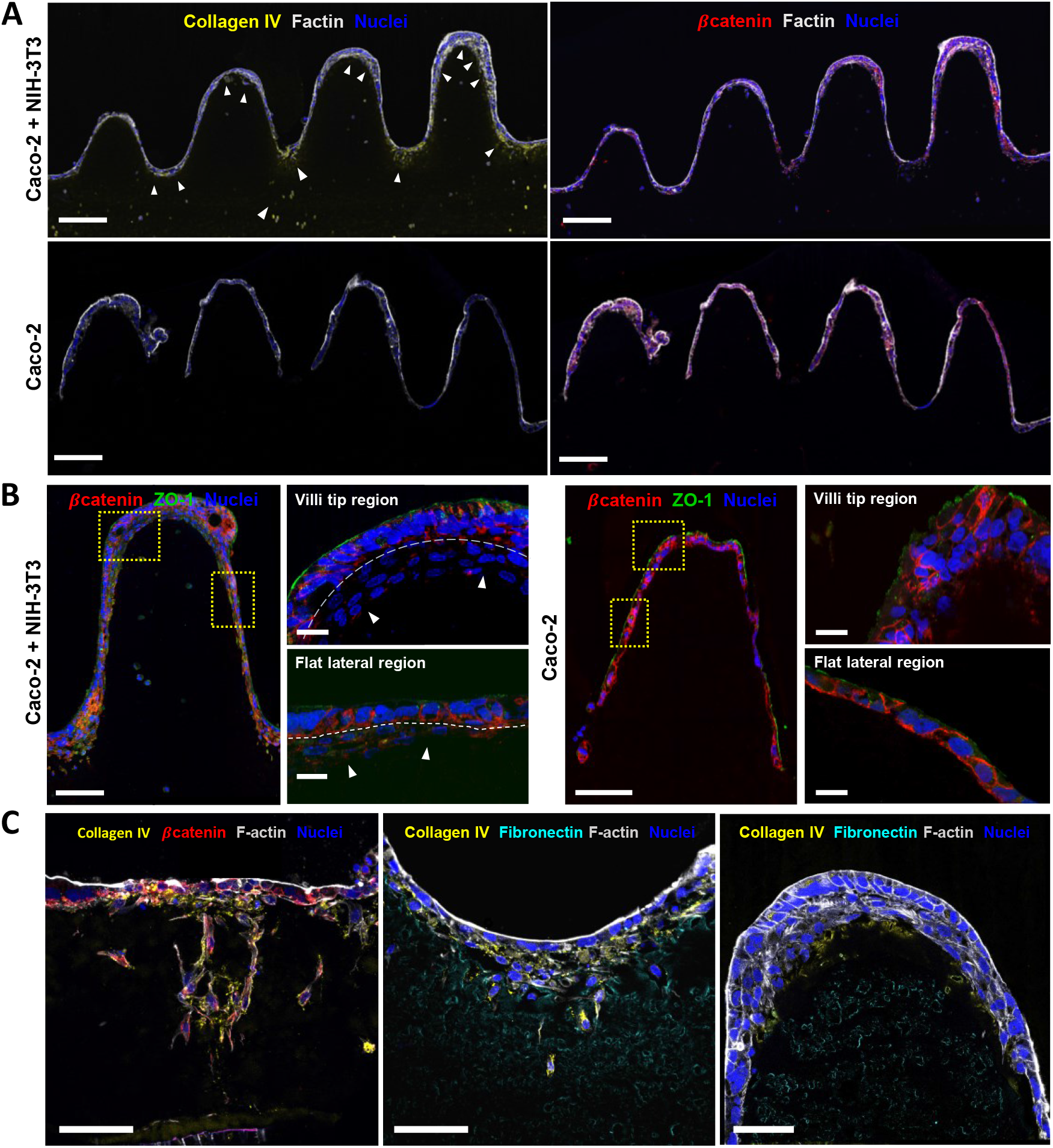
Confirmation of the 3D printed intestinal mucosa model. Immunostainings of the main markers for both the epithelial and the stromal compartments. (A) F-actin (grey) and nuclei (blue) showing cell distribution along the cross-sections of the 3D prints for Caco-2 and NIH-3T3 co-culture (left) and control (right) samples. Collagen IV (yellow, top) and β-catenin (red, bottom) were chosen as specific markers for fibroblasts and Caco-2 cells, respectively. Arrow tips show regions where fibroblasts migrated forming a basement below the epithelial monolayer. Scale bars = 200 μm. (B) β-catenin (red) and ZO-1 (green) show epithelial cells polarization at different regions of the microstructures for both sample types. Sale bars = 100 μm., and 20 μm inserts. (C) Collagen IV (yellow) and fibronectin (cyan) evidence the fibroblast activity within the gels, at different sample regions. Sale bars = 50 μm. All samples were fixed and stained after 21 days of culture.

Regarding the samples containing the fibroblasts, those were distinguishable from the epithelial cells because of their morphology (non-columnar or cuboidal) and their nuclear staining. Noteworthy, it was observed that after 21 days in culture the distribution of the fibroblasts along the sample was dramatically different depending on their position. Thus, while cell density within the bulk and the core of the villi was quite low, fibroblasts accumulated at the tips of the villi and the crypt regions (Figs. 8B, 8C). Such uneven fibroblast distribution along the crypt-villous axis has a striking resemblance to what has been previously reported in the field (8). Moreover, not only fibroblast distribution but also their morphology and appear as dependent on their specific location across the sample. Fibroblasts surrounded by hydrogel showed a roundish morphology. Meanwhile, fibroblasts closer to the epithelial layer exhibited a spread morphology with either a migratory phenotype towards the epithelium (Fig. 8C left) or an elongated shape below the villous tips and crypt regions (Fig. 8B middle and right). Recently, it has been demonstrated that the local curvature of cell culture substrates plays an important role in tissue architecture, thus emphasizing the relevance of including 3D structures to retrieve *in vivo* physiology *in vitro* (61–63). Noticeably, fibroblasts located at the crypt regions showed increased expression of collagen IV and fibronectin, which are significantly diminished from those located at the tips of the villi (Fig. 8C right), therefore suggesting different functionalities between those two cell populations. When in co-culture, fibroblasts show the capability of remodelling the matrix acting as basement membrane, thus providing a proper cellular microenvironment for the growth and migration or the epithelial cells, promoting the maturation of their junctions (10, 60, 64).

## Discussion

The use of light-based 3D printing techniques has grown exponentially in the past years, becoming one of the most promising fabrication strategies in the bioengineering field (19). Of particular interest is the combination of bioinks with cells to produce complex 3D functional living tissues (18, 19). However, the presence of living cells imposes specific requirements on the printing process to avoid cell stress and toxicity. Specifically, the nature and concentration of the photoinitiator employed, the printing time and dose, and the light source used should be carefully controlled (19).

Here we have presented a customized DLP-based bioprinting system based on visible-light photopolymerization suitable for the controlled printing of cell-laden bioinks with low macromer content. The apparatus was successfully adapted for printing highly transparent photosensitive polymers in reduced volumes, including temperature sensitive materials such as GelMA, and cell-laden bioinks, leading to high cell viability rates. Despite similar systems have been previously introduced (14, 65–67), it is still challenging to faithfully crosslink solutions of low macromer content (< 10% w/v), which are key to guarantee the survival of cells in soft tissues (46, 47). These polymeric solutions render hydrogels that possess variable swelling and mechanical properties in the range of those featured by soft tissues (< 40 kPa) (13, 67), which can be adapted on demand to better recreate the particular physicochemical properties of different extracellular matrix tissues, including cell adhesion and matrix remodeling capabilities, while having good mass transfer properties to allow cell viability for long time periods. By investigating the printing parameters suitable to produce well defined and mechanically stable printed constructs of GelMA - PEGDA polymer solutions, we could estimate the impact of the main printing parameters (printed layer thickness, printing time per layer, exposure intensity, CAD design characteristics) on the bioprinting process. It is worth to mention that these parameters are entangled in printing process and, while on the one hand make pose some difficulties to find the proper conditions, on the other hand, when using a suitable bioink, provide the system with an easy, straightforward methodology to fine tune the crosslinking density of the hydrogel. Additionally, the optical characteristics of the bioink were also adjusted by adding a photosensitive dye, tartrazine, to the pre-polymer mixture (30, 32). We found that by adjusting the content of tartrazine, it is possible to modify the gelation kinetics of the bioink and achieve a better confinement of the crosslinking reaction, thus, minimizing the undesired overexposure effects. With this approach, we can produce mechanically stable hydrogel prints of complex architectures, as it was proved by the successful printing of samples containing crypt-villous structures mimicking the small intestinal tissue. To do that, we employed as bioink a mixture of a natural derived polymer (GelMA) and a synthetic polymer (PEGDA). By controlling its photopolymerization, this polymeric solution has been proven to lead hydrogels than can be tuned to match the mechanical requirements of the target tissue (intestine) while providing at the same time cell adhesion and remodeling cues (GelMA), and mechanical long-term stability (PEGDA) (7, 13, 47, 68).

Finally, we proved the benefits of our bioprinting system by creating functional *in vitro* models of the intestinal mucosa in a reproducible and reliable manner. Multiple works have addressed the importance of accounting with the 3D architecture of the small intestinal tissue when addressing questions going from basic biology till drug absorption, drug-pathogen interactions, or disease modeling (6, 7, 68, 69). The relevance of including in the models not only the epithelial but also the stromal compartment is also well recognized but challenging to achieve in a reliable and efficient manner. In here we show that by employing a bioink based on GelMA-PEGDA polymers and an optimized bioprinting procedure we could successfully obtain samples that not only feature the crypt-villous architecture of the native tissue, but also include the relevant cell populations of the stromal and epithelial compartments and its functionality as tissue barrier, thus providing a unique *in vitro* model of the intestinal mucosa. This model could be cultured for at least 21 days and allowed to perform TEER measurements and histological cuts, thus emphasizing its potential applications in the field of *in vitro* tissue modeling. Our experiments demonstrated obvious effects of crosstalk between the different cell populations. TEER values revealed a boost in the growth of the epithelial monolayer on top of the hydrogels containing embedded fibroblasts, which has been mainly attributed to the paracrine signaling via hepatocyte growth factor (HGF) (57) and keratinocyte growth factor (KGF) (56), among others. Noteworthy, histological studies also revealed that close to the surface, fibroblasts showed a migratory phenotype towards the epithelium and the accumulation of secreted collagen IV forming a basement membrane to sustain the epithelial monolayer and thus remodeling the matrix, as already reported for the stromal tissue (8). They also showed a characteristic distribution of the fibroblasts within our 3D bioprinted model, which accumulate at the tips of the villi and the crypt regions (corresponding to regions with convex and concave surface topography), similar to what has been shown *in vivo* (59–62).

In summary, here we have proposed a light-based 3D bioprinting approach as a feasible alternative for developing *in vitro* cell culture models recapitulating the native microenvironment of the *in vivo* tissue, thus contributing on providing alternatives beyond the current state-of-the-art. Remarkably, the approach presented here does not have any inherent constraint to be scalable to the size of a conventional well-plate, thus providing a suitable alternative to include complex 3D models of tissues in standard assays.

## Materials and Methods

### Materials

Gelatin methacryloil (GelMA) was prepared following a method previously described (70, 71). Briefly, a 10% (w/v) gelatin solution was first obtained by dissolving gelatin from porcine skin type A (Sigma-Aldrich) in phosphate buffer saline (PBS; pH 7.4) (Gibco) at 50°C for 2 h under stirring conditions. Methacrylic anhydride (Sigma-Aldrich) at 1.25% v/v was added at a rate of 0.5 mL/min and left to react for 1 h while stirring. The solution was centrifuged at 1200 rpm for 3 min and the reaction was stopped by adding Milli-Q water. The resulting solution was dialyzed using 6-8 kDa molecular weight cut-off membranes (Spectra/por, Spectrumlabs) against Milli-Q water at 40°C, which was replaced every 4 h for 3 days. Then, after adjusting the pH to 7.4, the dialyzed products were frozen overnight at −80°C and lyophilized for 4-5 days (Freeze Dryer Alpha 1-4 LD Christ). The resulting product, GelMA, with a methacrylation degree of 47.5 ± 4% (determined by Trinitrobenzene sulfonate assay, data not shown) was stored at −20°C until further use.

A set of printing solutions (bioinks) was then prepared by mixing GelMA, poly(ethylene glycol) diacrylate (PEGDA) with a molecular weight of 4000 Da (Polysciences), visible-light lithium phenyl-2,4,6-trimethylbenzoylphosphinate (LAP) photoinitiator (TCI Europe), and tartrazine food dye (Acid Yellow 23, Sigma-Aldrich) at different w/v ratios and combinations using either Hank’s Balanced Salt Solution (HBSS) (Sigma-Aldrich) or high glucose Dulbecco’s Modified Eagle Medium (DMEM) without phenol red (Gibco) as dilution buffers.

### DLP-SLA-based bioprinting platform

A customized digital light processing stereolithography (DLP-SLA) 3D bioprinting system was built by modifying a commercially available, low cost Solus 3D printer (Junction3D), initially designed to print hard resins. As shown in Fig. 1A, the system consists of the following main components: a facing-down printing support coupled to a Z-axis motor, a resin vat with a transparent window, and a beam projector. A customized aluminum printing support (12 mm of diameter) and an aluminum vat (20 mm of inner diameter) were designed for printing small samples (between 3 to 10 mm in diameter) using reduced prepolymer volumes (< 2 mL). Aluminum was chosen for its good thermal conductivity and its lower oxygen permeability. A FEP film (Junction3D) of 150 μm in thickness was used to create a flexible transparent window at the bottom of the vat to allow the pattern transfer to the prepolymer solution. This creates, at the same time, an oxygen permeable window to tune the free radical photopolymerization reaction. Additionally, a flexible heater with a thermostat (TUTCO) was coupled to the system to keep the prepolymer solutions warmed at 37°C, preventing physical gelation, and allowing for the use of cell-laden polymers as bioinks. A full High-Definition 1080p resolution projector (Vivitek) was employed to crosslink the polymeric network using visible light. To avoid cell damage due to IR radiation exposure, a short pass heat protection filter (Schott) was added to the output of the projector. To form the 3D printed design, series of focused white and black patterns were projected onto the photocrosslinkable prepolymer solution through the transparent vat window, forming the 3D structure in the vertical direction, layer by layer (Fig. 1B). The optical power density applied for printing was set between 3.1 and 12.3 mW/cm^2^ (measured at the printing plane) through 5 predefined intensity levels in the projector (see Fig. S2) within the 320 to 640 nm wavelength spectral range. For the bioink compositions and the designs selected here, printing parameters were varied as following: layer thickness between 10 and 25 μm and exposure time per layer between 1 and 15 s.

### Bioprinting procedure

Printed designs in the shape of discs and the crypt-villus architecture of the small intestinal tissue were tested with the customized DLP-SLA platform. GelMA-LAP, PEGDA-LAP and GelMA-PEGDA-LAP were used as bioinks, always keeping the total macromer content to 8% w/v, according to previous findings (13). For printing, first PEGDA and GelMA polymers were dissolved separately in HBSS supplemented with 1% v/v Penicillin-Streptomycin (Sigma-Aldrich) at 65°C under stirring conditions for 2 h. Then, PEGDA solutions were filtered and mixed with the corresponding percentages of GelMA (when required) and LAP. To minimize undesired overexposure effects and have a finer control of the photocrosslinking process, the photoabsorber tartrazine was added to the mixtures. The final prepolymer solutions were kept at 37°C for about 30 min before use. Then, the working prepolymer solution was pipetted into the vat of the DLP-SLA system, which was previously warmed at 37°C. The design to be printed was programmed in CAD and the printing parameters were adjusted onto the software accordingly. The print was created layer by layer from the bottom, on top of a printing support. To ensure the proper detachment of the samples from the printing support while avoiding damage, hydrogels were printed either on top of 12 mm diameter glass coverslips or 10 mm diameter Tracketch^®^ polyethylene terephthalate (PET) membranes with 5 μm pore size (Sabeu GmbH & Co). Both substrates were previously silanized to improve the attachment of the printed hydrogels (6). Once the printing process was finished, the entire pieces (supports plus hydrogels) were rinsed in warmed PBS at 37°C to remove unreacted polymer before being transferred to cell culture well plates and/or Transwell^®^ inserts and stored in PBS at 4°C until further use.

The influence of the relevant printing parameters (photoabsorber content, exposure time per printed layer and the printed layer thickness on the bioprinting outcome was checked using CAD models, which include a disc-like base with an array of bullet-shape pillars, mimicking the intestinal villi. The bioink employed was composed of 5% w/v GelMA, 3% w/v PEGDA and 0.4% w/v LAP and the projector power density was set to 12.3 mW/cm2. Each printing parameter (see Table 1) was evaluated individually in a systematic manner, and the resulting prints were analyzed right just after fabrication to avoid possible distortions due to hydrogel swelling.

### Swelling behaviour of the 3D printed hydrogels

The swelling behaviour of the printed hydrogels was assessed by monitoring the variations in sample dimensions in both, radial and transversal directions. Disc-like samples of 6 mm in diameter and 0.5 mm in height were printed onto glass coverslips using a power density of 12.3 mW/cm^2^, a layer thickness of 13 μm and 4 s of exposure time per printed layer. Right after photopolymerization, hydrogels were washed out using warmed HBSS and wiped with a KimWipe tissue to remove any excess of liquid. Pictures from top and lateral sides were taken using a stereo microscope (Olympus, SZX2-ILLB) and processed using ImageJ software (http://imagej.nih.gov/ij, NIH) to have the initial diameter and height values, *D_0_* and *H_0_*, respectively. After that, samples were kept submerged in HBSS with 1% v/v Penicillin/Streptomycin at 37°C to induce swelling until reaching equilibrium, exchanging the HBSS buffer solution every day. To monitor the process, pictures were taken at appointed times up to 96 hours. The swelling ratios of the printed hydrogels were determined using the following expressions,

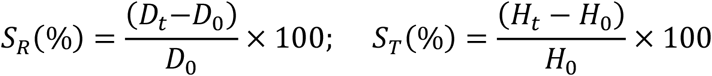

where *D_t_* and *H_t_* are the sample diameter and height values at time ‘t’, and SR and ST represent the swelling ratios in both radial and transversal directions, respectively. The volumetric swelling ratio, SV (%), was also evaluated using the expression,

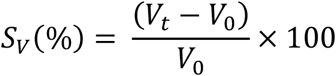

were *V_0_* is the initial sample volume, and *V_t_* is the sample volume at time ‘t’.

### Characterization of the mechanical properties of the 3D printed hydrogels

To determine the bulk mechanical properties and flow characteristics of the printed hydrogels, rheological measurements were performed. First, disc-like samples of 8 mm in diameter and 0.5 mm in height were printed using the selected bioink formulation, 5% w/v GelMA, 3% w/v PEGDA and 0.4% w/v LAP. As controls, prepolymer solutions containing, 8% w/v PEGDA + 0.4% w/v LAP, and 8% w/v GelMA + 0.4% w/v LAP were also included in these experiments. Samples were allowed to reach equilibrium swelling by being submerged in HBSS at 4°C for 5 days. As sample dimensions changed after swelling, they were punched to obtain 8 mm diameter samples for testing. Rheological tests were performed on a MCR302-PP08 rheometer (Anton Paar) equipped with parallel sandblasted plates of 8 mm in diameter. Measurements were performed using a sinusoidal signal and applying a sweep to the amplitude of shear strain between 0.01% and 500% while keeping constant the angular frequency to 10^-1^ s. The running temperature was maintained at 23°C throughout the measurements. Values of both storage (G’) and loss (G’’) moduli were obtained as a function of the strain, from which the elastic component of the moduli, *E*, was derived assuming a value for the Poisson’s ratio of 0.5.

### Characterization of the diffusivity properties of the 3D printed hydrogels

A drug diffusion study was performed to experimentally determine the mesh size of the printed hydrogels, since other methods, such as the one based on the Flory-Rehner theory, are not applicable when working with hydrogel co-networks (72). To do so, the diffusion profiles of dextran fluorescent molecules of different molecular weights passing through the hydrogels were analyzed. Disc-like samples of 6.5 mm in diameter and 0.5 mm in height were printed using pre-polymer solutions containing 5% w/v GelMA, 3% w/v PEGDA, 0.4% w/v LAP and 0.025% w/v of tartrazine in HBSS supplemented with 1% v/v Penicillin-Streptomycin. After printing, hydrogels were mounted on Transwell^®^ inserts using double-sided pressure sensitive adhesive (PSA) rings (Adhesive research) and left in standard cell culture conditions (37 °C and 5% CO_2_) to reach equilibrium swelling. Dextran molecules of 4 kDa (FITC-Dextran), 70 kDa (Rhodamine-Dextran), 150 kDa (FITC-Dextran), 500 kDa (FITC-Dextran) and 2000 kDa (FITC-Dextran) (all from Sigma-Aldrich) were used separately at a concentration of 0.5 mg/mL in PBS. Then, 200 μL of the pre-warmed dextran solutions were loaded in the apical compartments while adding 600 μL of PBS to the basolateral ones. The plates were then incubated at 37 °C and at regular time intervals, 50 μL were withdrawn from the basolateral compartments and replaced with warmed PBS. Collected samples were transferred to black 96-well plates and the FITC or Rhodamine fluorescence was measured using an Infinite M200 PRO Multimode microplate reader (Tecan), at excitation/emission wavelengths of 490/525 nm and 540/625 nm, respectively. The changing concentration of dextrans over time was determined using standard calibration curves and the diffusion coefficients (D) were calculated following the method form (55, 73)

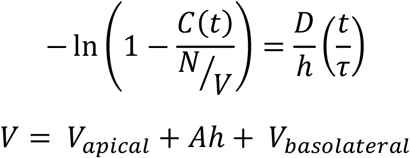

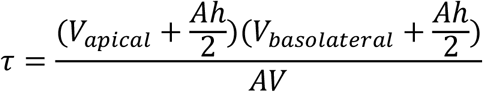

where *C*(*t*) is the dextran concentration in the basolateral chamber over time *t, N* is the total dextran mass, *h* is the height of the hydrogels, *V* is the total volume and *A* is the area in which the diffusion occurs.

### Cell culture

NIH-3T3 fibroblasts (ATCC^®^ CRL-1658™) from passages 12 to 17 were used for printing cell-laden hydrogels to mimic the cell population within the stromal compartment of the intestinal mucosa. Fibroblasts were grown, expanded, and maintained at 37°C and 5% CO_2_ in 175 cm^2^ culture flasks using high glucose DMEM (Gibco, ThermoFisher), supplemented with 10% v/v fetal bovine serum (FBS) (Gibco, ThermoFisher) and 1% v/v Penicillin-Streptomycin (Sigma-Aldrich). Cells were passaged twice a week, exchanging the medium every other day.

Caco-2 cells (ATCC^®^ HTB-37TM) from passages 83 to 87 were seeded on top of the printed hydrogels (with and without embedded cells) to mimic the epithelial layer of the small intestine. Caco-2 were grown, expanded, and maintained in 75 cm^2^ culture flasks in high glucose DMEM (Gibco, ThermoFisher), supplemented with 10% v/v fetal bovine serum (FBS) (Gibco, ThermoFisher), 1% v/v Penicillin/Streptomycin (Sigma-Aldrich) and 1% v/v of non-essential amino acids (NEA) (Gibco, ThermoFisher). Cells were maintained at 37°C and 5% CO_2_, changing medium every other day and passaged weekly.

### Fabrication and characterization of cell-laden, villous-like hydrogels: mimicking the small intestinal tissue

Prior to start with the fabrication of cell-laden samples, some prints were performed with the already optimized bioink to optimize the CAD models, leading to a variety of villus and crypt-like structures targeting the dimensions of the *in vivo* intestinal tissue (46). A prepolymer solution containing 5% w/v GelMA, 3% w/v PEGDA, 0.4% w/v LAP and 0.025% w/v of tartrazine in HBSS supplemented with 1% v/v Penicillin-Streptomycin, was used. Samples were fabricated on silanized PET membranes by using a printing optical power density of 12.3 mW/cm^2^. After printing, samples were washed out with warm PBS to remove the unreacted polymer. Bright field pictures from top and lateral view were taken using a stereo microscope (Olympus, SZX2-ILLB) and processed with ImageJ software (http://imagej.nih.gov/ij, NIH) to obtain the printed dimensions of the different elements in the CAD models. The base thickness, the villus height and crypt diameter and depth were obtained from samples cross-sections, after cutting them with a blade (see Figure 4). All these values were then compared to the ones on the CAD designs and adjusted, accordingly, to reach a final model, with the following dimensions: villi diameter of 300 μm and 700 μm in height, crypts diameter of 250 μm and 150 μm in depth, and base diameter of 6.1 μm and of 300 μm in height. Both the villous and crypt interdistances were set to 600 μm.

Cell-laden hydrogels with the villous morphology of the small intestinal tissue were then bioprinted to mimic the stromal compartment of the intestinal mucosa. NIH-3T3 fibroblasts were trypsinised from the cell culture flasks and re-suspended in the prepolymer solutions to get bioinks with a cell density of 7.5·10^6^ cells/mL. Then, bioinks were immediately placed in the vat previously warmed at 37°C and the printing process started. To maintain the cell suspension homogeneously distributed through the vat volume along the printing time, the pre-polymer solution was stirred by gently pipetting. Samples were fabricated on silanized PET membranes by using a printing optical power density of 12.3 mW/cm^2^. After the printing process, samples were washed out with warm cell culture medium supplemented with 1% of Penicillin-Streptomycin to remove the unreacted polymer. Then, the bioprinted cell-laden hydrogels were attached to Transwell^®^ inserts by PSA rings. Finally, Caco-2 cells were seeded on top of the samples at a density of 2.5·105 cells/cm2 to mimic the intestinal epithelial layer. The samples were cultured for 21 days in an incubator at 37°C and 5% CO_2_, replacing medium every other day.

### Viability of cells embedded onto the 3D printed hydrogels

Fibroblast viability within the bioprinted cell-laden hydrogels was investigated on hydrogels with a simplified geometry (3 x 3 squared grid 500 μm thick and including an array of 500 μm width squared posts, Fig. 6B). For this purpose, it was employed a calcein-AM/ethidium homodimer Live/Dead kit (Invitrogen), 24 h and 7 days after NIH-3T3 cell encapsulation. Hoechst was used as live staining for the nuclei. A confocal laser scanning microscope (LSM 800, Zeiss) was used for imaging and a manual cell counter plugin in ImageJ software (http://imagej.nih.gov/ij, NIH) was employed for image processing and cell viability quantification. Data was plotted using GraphPad Prism 9 software.

### Histological and immunofluorescence analysis

After 21 days in culture, the morphology and phenotype of both Caco-2 and NIH-3T3 cells present in the bioprinted hydrogels featuring the intestinal villi and crypts were studied by immunostaining. Histological cuts were performed for this purpose. To preserve the morphology of the villous and crypt-like features on the soft hydrogels, samples were embedded following a protocol developed by our (58) group. Briefly, after fixation with 10% neutral buffered formalin solution (Sigma-Aldrich) at 4°C for 1 h, samples were submerged in a prepolymer solution containing 10% w/v of PEGDA 575 kDa (Sigma-Aldrich) and 1% w/v of 2-Hydroxy-4’-(2-hydroxyethoxy)-2-methylpropiophenone (Irgacure D-2959) photoinitiator (Sigma-Aldrich) in phosphate buffered saline (PBS) (ThermoFisher) and kept overnight at 4 °C. Then, samples were placed within poly(dimethyl siloxane) (PDMS) (Dow corning) round pools of 12 mm in diameter and 2 mm in thickness attached to a plastic support, which was filled with PEGDA 575 kDa prepolymer solution. The construct was then irradiated using UV light at 365 nm wavelength in a MJBA mask aligner (SUSS MicroTech). To ensure the formation of a homogeneous block, samples were irradiated twice for 100 s at 25 mW/cm2 of power density, first from the top and later from the bottom, by flipping the sample downwards. An additional first exposure of 40 s was required to form a support base to keep the plastic support in place. After UV exposure, unreacted polymer and photoinitiator were washed out with PBS. Then, samples on PEGDA blocks were immersed overnight in 30% sucrose solution (Sigma-Aldrich) at 4°C for cryoprotection, further embedded in OCT (Tissue-Tek^®^ O.C.T. Compound, Sakura^®^ Finetek) and stored at −80 °C for at least 12 h. Finally, they were cut with a cryostat (Leica CM195) and histological cross-sections sections of ~7 μm thickness were recovered, attached onto glass coverslips, air dried and stored at −80°C until use.

For immunostaining, samples on glass coverslips were left to unfreeze at room temperature (RT) and washed carefully with PBS. Then, cells were permeabilized with 0.5% Triton-X (Sigma-Aldrich) at 4°C for 2 h and blocked with 1% bovine serum albumin (Sigma-Aldrich), 3% donkey serum (Millipore), and 0.3% Triton-X at 4°C for 2 h. Drops of 50 μL containing primary antibodies against β-catenin (Abcam) (1 μg/mL), collagen IV (Biorad) (1.6 μg/ mL), fibronectin (Santa Cruz) (2 μg/ mL) and Ki67 (BD Pharmingen) (2.5 μg/ mL) were placed on top of the sections and incubated overnight in a moisture chamber at 4°C under shaking conditions, and covered with Parafilm^®^ (Sigma-Aldrich) to prevent drying. After several washing steps with PBS, samples were incubated with the secondary antibodies anti-goat Alexa 647, and anti-rabbit Alexa 488 or antimouse Alexa 488 (Invitrogen, ThermoFisher) (4 μg/ mL) together with Rhodamine-Phalloidin (Cytoskeleton) (0.07 μM) for 2 h at 4°C, and finally, were incubated with DAPI (5 μg/·mL) for 30 min. In the case of anti ZO-1 staining (ThermoFisher) (2.5 μg/°mL), an antigen retrieval treatment was performed prior sample permeabilization to enhance its signal. Briefly, samples were boiled for 10 min after unfreezing in citrate buffer solution (10 mM citrate and 0.05 % v/v of Tween20 in MilliQ water, previously adjusting the pH at 6), controlling bubble formation. Then, the staining procedure continued as for the other markers. After immunostaining, samples were covered by thin glass coverslips with a drop of Fluoromount G (Southern Biotech) and were imaged using a confocal laser-scanning microscope (LSM 800, Zeiss).

### Epithelial barrier integrity measurements by monitoring the transepithelial electrical resistance (TEER)

The growth and integrity of the epithelial barrier formed on top of the villous-like printed hydrogels was assessed by monitoring the transepithelial electrical resistance (TEER) between the apical and basolateral compartments defined by the Transwell^®^ inserts. This was tracked three times per week using an EVOM2 Epithelial voltohmmeter with an STX3 electrode (World precision Instruments) for a total culture time of 21 days. The resistance values measured were corrected by subtracting the resistance of the porous PET membranes of the Transwell^®^ inserts and the resistance of the cell-free and cell-laden villous-like hydrogels. TEER values were normalized by the total surface area of the epithelial monolayers, taking into account the morphology of the printed features (6).

### Statistics and data analysis

Graphpad Prism 8 software was used for data treatment and analysis. The data in graphs are presented as mean values with standard deviation (SD). In the case of normal distributions, statistical significance was interrogated by means of one-way ANOVA test. Turkey’s test was also performed when indicated in the figure captions. Values of p < 0.05 were used to consider differences as statistically significant.

## Acknowledgments

Authors gratefully acknowledge Mr. Fernando Lucena, from Anton Paar company, for his help and advice during the rheology measurements.

## Funding

Funding for this project was provided by:

European Union Horizon 2020 ERC grant (agreement no. 647863 - COMIET)

European Union Horizon 2020 ERC-PoC grant (agreement no. 899906 – GUT3DPLATE)

European Union Horizon 2020 FET-Open grant (agreement no. 828931 – BRIGHTER),

Spanish Ministry of Science and Innovation, Severo Ochoa Programme for Centres of Excellence in R&D in Spain (2016–2019)

Spanish Ministry of Science and Innovation, Juan de la Cierva programme (grant IJC2019-040289-I), NT.

The results presented here only reflect the views of the authors; the European Commission is not responsible for any use that may be made of the information it contains.

## Author contributions

CRediT author statement

Conceptualization: EM, NT

Methodology: NT, JZ

Validation: JZ, NT, EA

Investigation: NT, JZ, EA, MG, AC

Visualization: NT, MG

Supervision: EM, NT

Writing—original draft: NT, MG, EM

Writing—review & editing: NT, EM

## Competing interests

All authors declare they have no competing interests.

Data and materials availability: All data needed to evaluate the conclusions in the paper are present in the paper and/or the Supplementary Materials.

## Supplementary Materials

Supplementary materials are available in a separate file.

