## Supplementary material for "A simple DLP-bioprinting strategy produces cell-laden crypt-villous structures for an advanced 3D gut model"

**This PDF file includes:**

Figs. S1 to S6

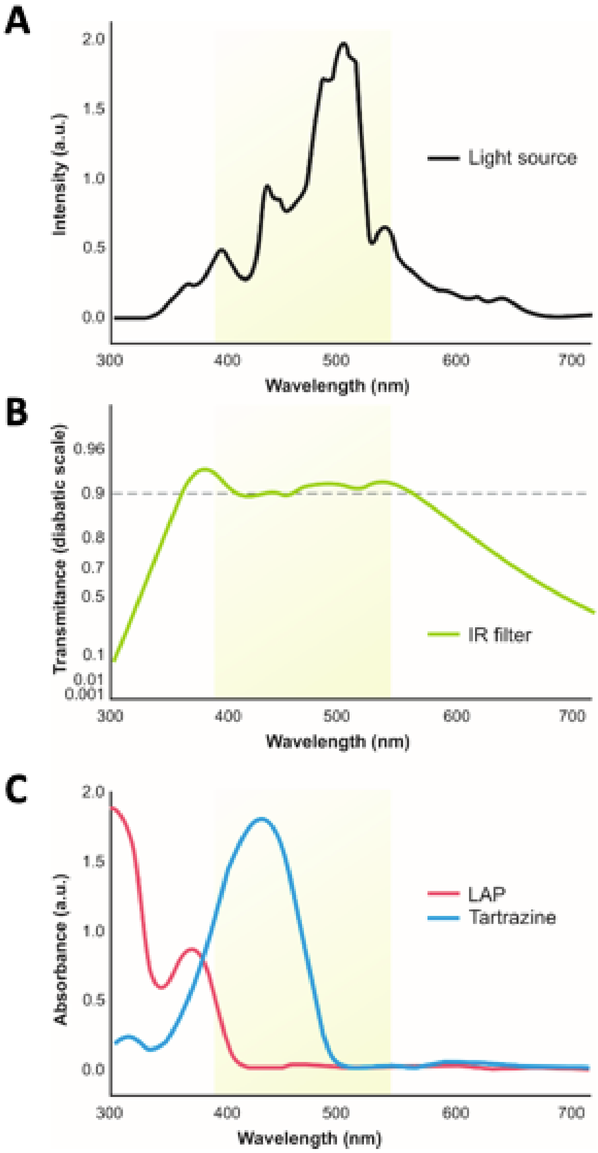


**Fig. S1.**

**Spectral analysis of the different components of the system:** (A) Spectrum of the visible light emitted by the projector beam. (B) Transmittance of the IR-filter; (C) UV-visible absorption spectra of the LAP photoinitiator and the tartrazine dye.

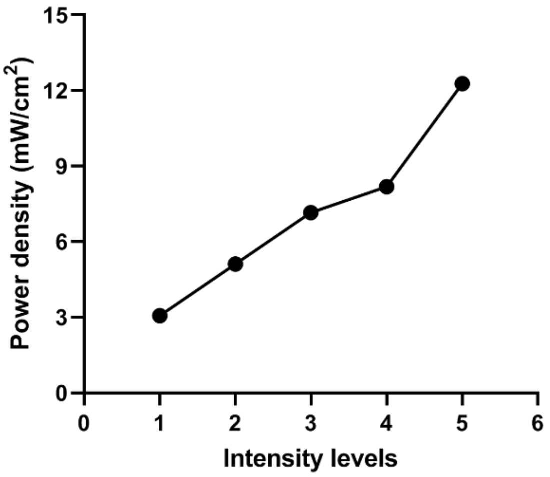


**Fig. S2.**

**Optical power density** of the 3D printing setup as function of the light intensity levels of the projector, measured at the printing plane.

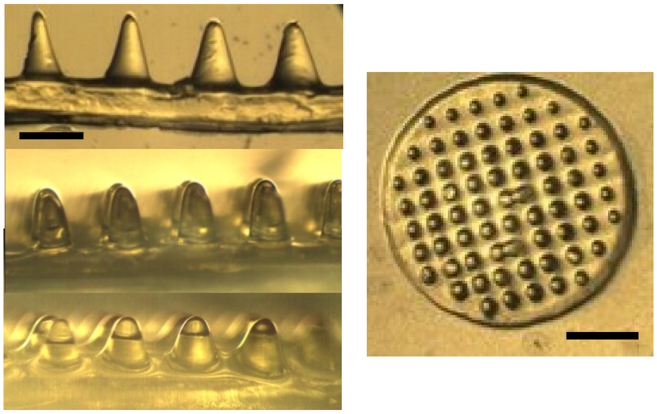

**Fig. S3.**

**Tuneable morphology of the printed structures.** Using the same bioink composition and printing parameters, villous-like structures with different height and diameter can be obtained only varying the CAD design. Scale bars = 400 µm (lateral views), 1.5 mm (top view).

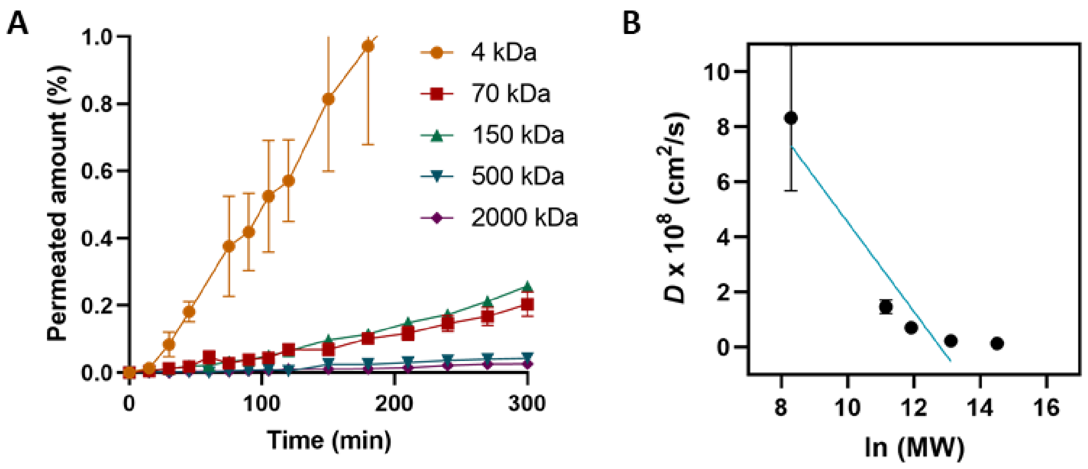


**Fig. S4. Diffusive properties of the 3D bioprinted hydrogels.**

Graphs showing (**A**) the amount of permeated dextran molecules through 3D printed disc-like hydrogels, and (**B**) the diffusion coefficients versus the ln of the dextran's molecular weight, from which the size exclusion limit was calculated. In both cases, dextran molecules of 4 kDa (FD4), 70 kDa (FD70), 150 kDa, 500 kDa (FD500), and 2000 kDa (FD2000) were tested on hydrogels printed using 5% w/v GelMA, 3 % w/v PEGDA, 0.4 % w/v LAP, and 0.025% v/v tartrazine bioink and the following optimized printing parameters: 13 µm of layer thickness and 5 s of exposure time, with an exposure dose of 61.5 mJ/cm$^2$ per printed layer. Values are shown as mean ± SD (min n=3).

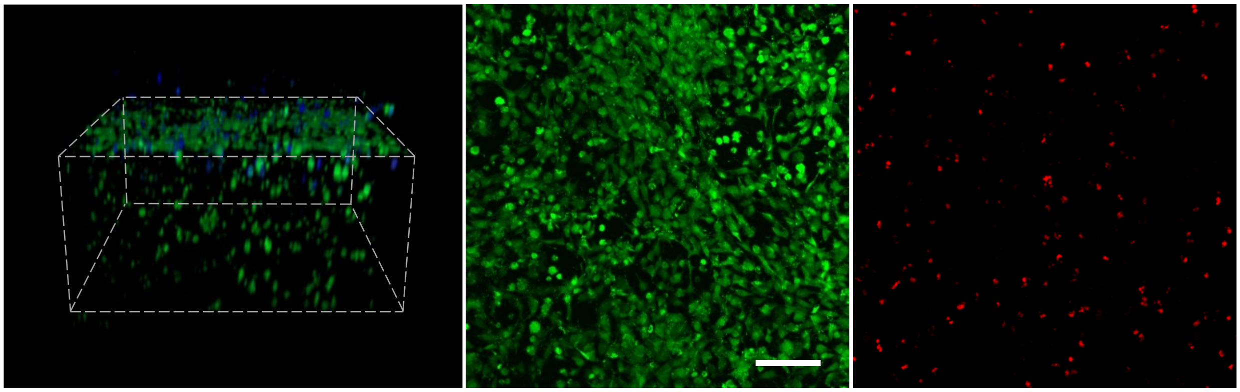

**Fig. S5.**

**NIH-3T3 network on top hydrogel layers.** 10 days post-encapsulation, NIH-3T3 fibroblasts formed a dense network on the top layers of 3D bioprinted hydrogels. Scale bar = 100 µm.

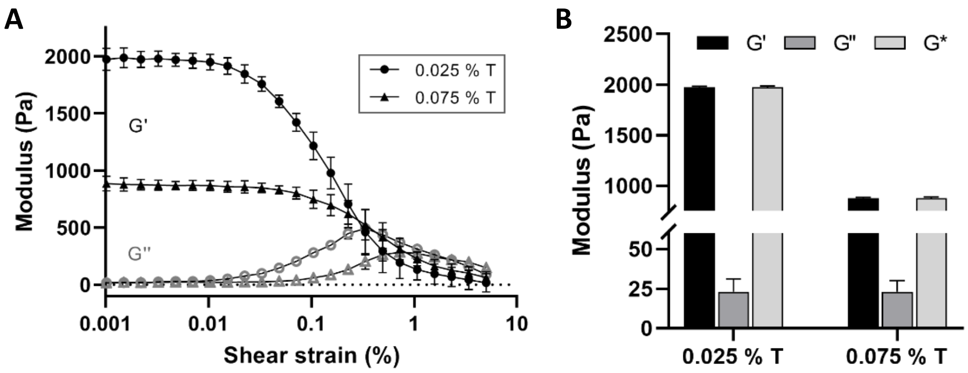


**Fig. S6.**

**Impact of increasing the tartrazine content on the mechanical properties of the printed hydrogels.** (**A**) Rheological curves showing G’ and G’’ moduli and (**B**) comparison of complex, storage and loss moduli, for hydrogels containing both 0.025% and 0.075% w/v concentration of tartrazine.
